# PhyMorphS: A Physics-Guided Framework for Mesh-Based Morphological Synthesis of White Matter Microstructures

**DOI:** 10.64898/2026.11.09.765314

**Authors:** Ruiqi Cui, Mahsa Amirrashedi, Mario Corral-Bolaños, Jean-Philippe Thiran, J. Andreas Bærentzen, Tim B. Dyrby

## Abstract

Numerical phantoms provide controlled 3D representations of brain tissue microstructure for developing and evaluating biophysical models of diffusion-weighted Magnetic Resonance Imaging (dMRI). Existing phantom synthesizers employ different geometric primitives and optimization strategies to generate densely packed substrates with diverse axonal morphologies. However, direct deformation of axonal surfaces under mechanical constraints remains less explored. We propose PhyMorphS, a mesh-based framework in which axonal morphology emerges through physics-guided growth and interactions with the local microstructural environment, without constraining cross-sections to predefined symmetric shapes. In PhyMorphS, axons, myelin sheaths, and cell bodies are represented by triangle meshes, while a mass-spring system and collision-resolution scheme guide iterative axonal growth under mechanical and packing constraints. This enables irregular, asymmetric, and non-convex cross-sections, longitudinal diameter variation, and user-defined morphological alterations such as axonal beading and myelin layer decompaction. We evaluate PhyMorphS against large-scale X-ray nano-holotomography segmentations of real 3D axons from four white matter regions with distinct fiber organizations. The synthesized axons reproduce key region-specific morphological characteristics and achieve high packing densities, while the parallelized implementation scales approximately linearly with axon count in the generation voxel. By reproducing the geometric variability observed in real WM tissue, PhyMorphS provides controlled substrates for investigating how realistic morphology influences simulated dMRI signals, testing the geometric assumptions of biophysical microstructure models, and generating diverse paired microstructure-signal datasets for training machine-learning models.

**Graphical Abstract:** 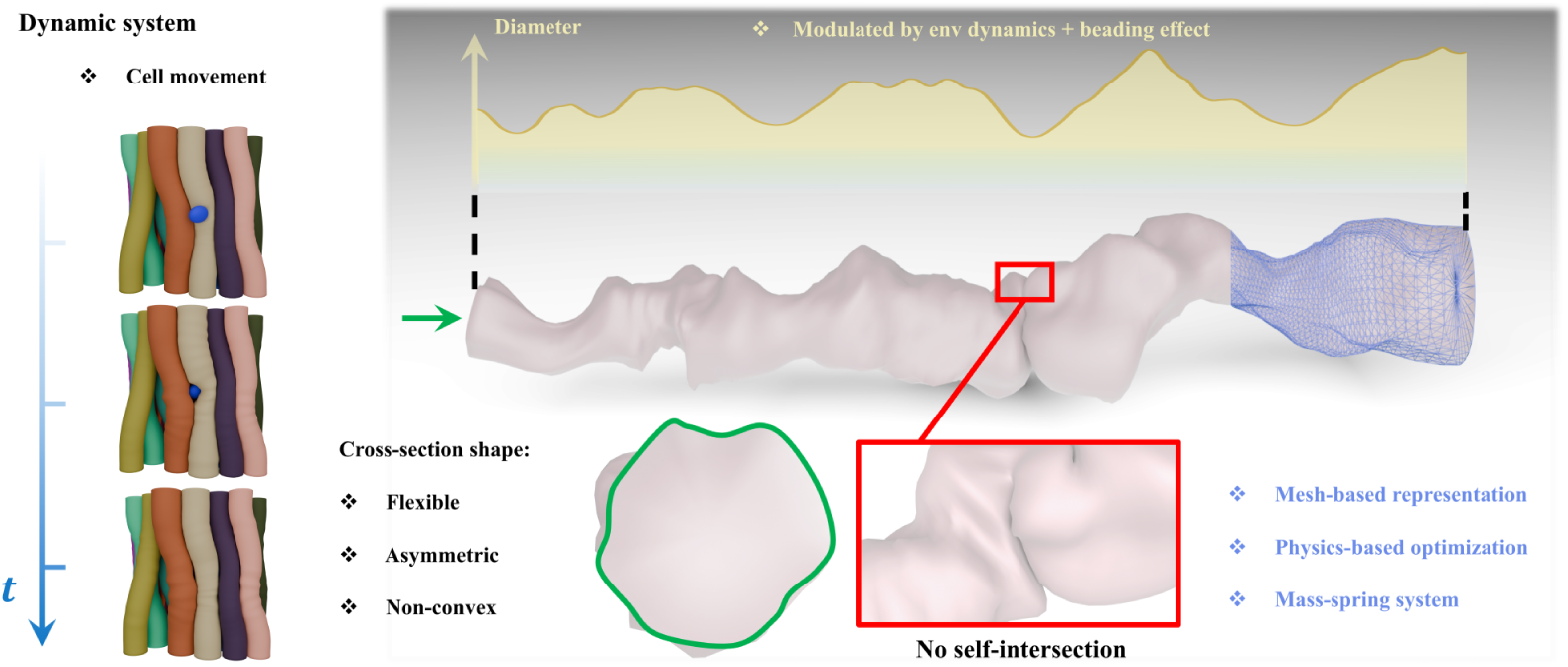

**Highlights:**

- PhyMorphS is a physics-based synthesizer that generates realistic brain WM microstructures.
- Synthesized axons reproduce XNH-derived diameter, cross-sectional shape, and tortuosity.
- Physics-guided deformation enables plausible dynamics and morphological alterations.
- Extensive evaluations against large-scale real data validate PhyMorphS realism.

## 1. Introduction

Numerical phantoms of brain white matter (WM) provide controlled three-dimensional (3D) representations of tissue for simulating diffusion-weighted magnetic resonance imaging (dMRI) signals and developing and evaluating biophysical models (Dyrby et al., 2018; Hall and Alexander, 2009; Rafael-Patino et al., 2020; Fieremans et al., 2008). These models infer microstructural properties from water diffusion, often under simplifying assumptions about tissue geometry (Alexander et al., 2019; Assaf and Basser, 2005; Assaf et al., 2008; Zhang et al., 2012). Phantoms with known geometry allow these assumptions to be tested and the effects of specific morphological features to be examined. A central challenge is to reproduce the diversity of axonal shapes and their organization within the surrounding tissue, which vary across anatomical regions and spatial scales (Andersson et al., 2020; Kjer et al., 2025).

3D electron microscopy (EM) and X-ray nano-holotomography (XNH) have revealed irregular axonal cross-sections, longitudinal diameter variations, and tortuous trajectories (Abdollahzadeh et al., 2019; Amirrashedi et al., 2025; Andersson et al., 2020). These features influence diffusion measurements and their interpretation, including estimates of axon diameter and the effects of susceptibility-induced internal gradients (Andersson et al., 2020; Lee et al., 2020a; Nilsson et al., 2012; Lee et al., 2020b; Winther et al., 2024a). Phantoms that reproduce these features therefore enable controlled investigation of how geometric assumptions affect dMRI interpretation.

Existing synthesizers, including MEDUSA (Ginsburger et al., 2019), CATER-Pillar (Nguyen-Duc et al., 2025), CACTUS (Villarreal-Haro et al., 2023), WMG (Winther et al., 2024b), and ConFiG (Callaghan et al., 2020), combine geometric packing and growth rules to generate densely packed substrates. Representations based on simple geometric primitives provide computational efficiency but can restrict local cross-sectional shape. For example, MEDUSA(Ginsburger et al., 2019) represents axons as chains of spheres, restricting local cross-sections to circular shapes. WMG (Winther et al., 2024b) relaxes this restriction by using ellipsoids, allowing eccentricity while retaining elliptical cross-sections. ConFiG (Callaghan et al., 2020) and CACTUS (Villarreal-Haro et al., 2023) generate flexible axonal morphologies through surface construction and voxel-based radial growth with interactions between neighboring fibers, respectively. These approaches can produce irregular cross-sections and complex axonal trajectories. PhyMorphS differs in explicitly modeling surface elasticity and internal mechanical support through a mass-spring system coupled to deformable surface meshes, allowing local surface deformation to be governed by physical constraints to achieve higher realism.

Previous synthesizers have been evaluated using a range of morphological measures and anatomical references (Callaghan et al., 2020; Ginsburger et al., 2019; Nguyen-Duc et al., 2025; Villarreal-Haro et al., 2023; Winther et al., 2024b). However, comprehensive comparisons of cross-sectional morphology and regional variability across multiple WM regions remain limited. Broader evaluations are therefore needed to assess how well synthesized phantoms reproduce the diversity of axonal morphology observed in real tissue.

In this paper, we propose PhyMorphS, a physics-guided framework for mesh-based synthesis of WM microstructure. Building on our preliminary mass-spring formulation (Cui et al., 2024), PhyMorphS combines deformable triangle meshes with a mass-spring system to guide axonal growth and deformation under mechanical and packing constraints. Collision handling accounts for interactions among axons and cell bodies, allowing asymmetric and non-convex cross-sections to emerge. Beyond the preliminary work, we introduce data-driven initialization from regional orientation and diameter distributions, revised spring connectivity and growth constraints for robust synthesis under dense packing, and parallelized optimization and collision processing. The framework also supports myelin representation and controlled morphological alterations, including axonal beading and myelin layer decompaction.

We further extend the evaluation through quantitative comparisons with high-resolution XNH segmentations containing thousands of axons from four WM regions with distinct anatomical organization and complexity: aligned fibers in the splenium and midbody of the corpus callosum, crossing fibers in the centrum semiovale, and U-fibers beneath the cortex (Amirrashedi et al., 2025). We assess dense packing using a target fiber volume fraction (FVF) of 80%. We combine quantitative comparisons of axon diameter distributions, longitudinal diameter variation, cross-sectional eccentricity and convexity, and tortuosity with qualitative assessment to examine how well PhyMorphS captures anatomical variability across these regions.

## 2. Methods

### 2.1. Modeled tissue anatomy

PhyMorphS represents the WM microenvironment using axons, optional myelin sheaths, and glial cell bodies. Throughout this work, a *voxel* denotes the bounded three-dimensional domain within which the microstructural substrate is initialized and synthesized. Axonal and somatic interiors define the intracellular compartments, while the volume outside the axons, myelin sheaths, and cell bodies defines the extracellular compartment. Axons are modeled as deformable tubular structures with spatially varying trajectories, cross-sectional shapes, and diameters, reflecting anatomical observations (Andersson et al., 2020). Axonal diameters typically range from sub-micrometer values to several micrometers, depending on the brain region and fiber population (Alexander et al., 2019). We treat the axonal interior as homogeneous and do not explicitly resolve cytoskeletal structures (Kevenaar and Hoogenraad, 2015).

For myelinated axons, the sheath consists of closely packed membrane layers, or lamellae, separated by nanometer-scale water-filled spaces (Canales-Rodríguez et al., 2024). We approximate this structure using concentric mesh layers. Relative sheath thickness is specified by the g-ratio, defined as the inner axonal diameter divided by the outer fiber diameter, including myelin (Rushton, 1951; Skoven et al., 2025). The framework also supports axonal beading, modeled as alternating local swellings and constrictions (Budde and Frank, 2010; Lee et al., 2020b), and myelin layer decompaction, modeled as increased spacing between layers.

Cell bodies are modeled as ellipsoidal structures with user-defined sizes based on anatomical measurements (Andersson et al., 2020). Unlike axons, whose diameter can vary along their trajectories, cell bodies are modeled with a relatively constant size. Depending on the cell type, some glial cells, such as oligodendrocytes, remain relatively fixed in position, whereas others, including astrocytes and microglia, may move within the tissue environment (Tønnesen et al., 2018). Many glial cells possess thin processes extending from the cell body, however, these structures are not explicitly represented in the current framework. Instead, we simplify cells as rigid bodies represented solely by their soma, as observed in XNH imaging, capturing their dominant spatial influence on the organization of the tissue microenvironment while maintaining computational efficiency.

### 2.2. Voxel configuration and mesh initialization

Synthesis starts from a JSON configuration file containing one entry for each axon or cell body (Fig. 1.F). Each configuration defines a voxel to be synthesized. Table 1 summarizes the inputs. Each cell body is defined by its center *p_c_* and three semi-axis vectors **a**, **b**, and **c**, whose directions and lengths determine its orientation and size. Geometric inputs can be specified from anatomical measurements (Andersson et al., 2020).

**Figure 1:**
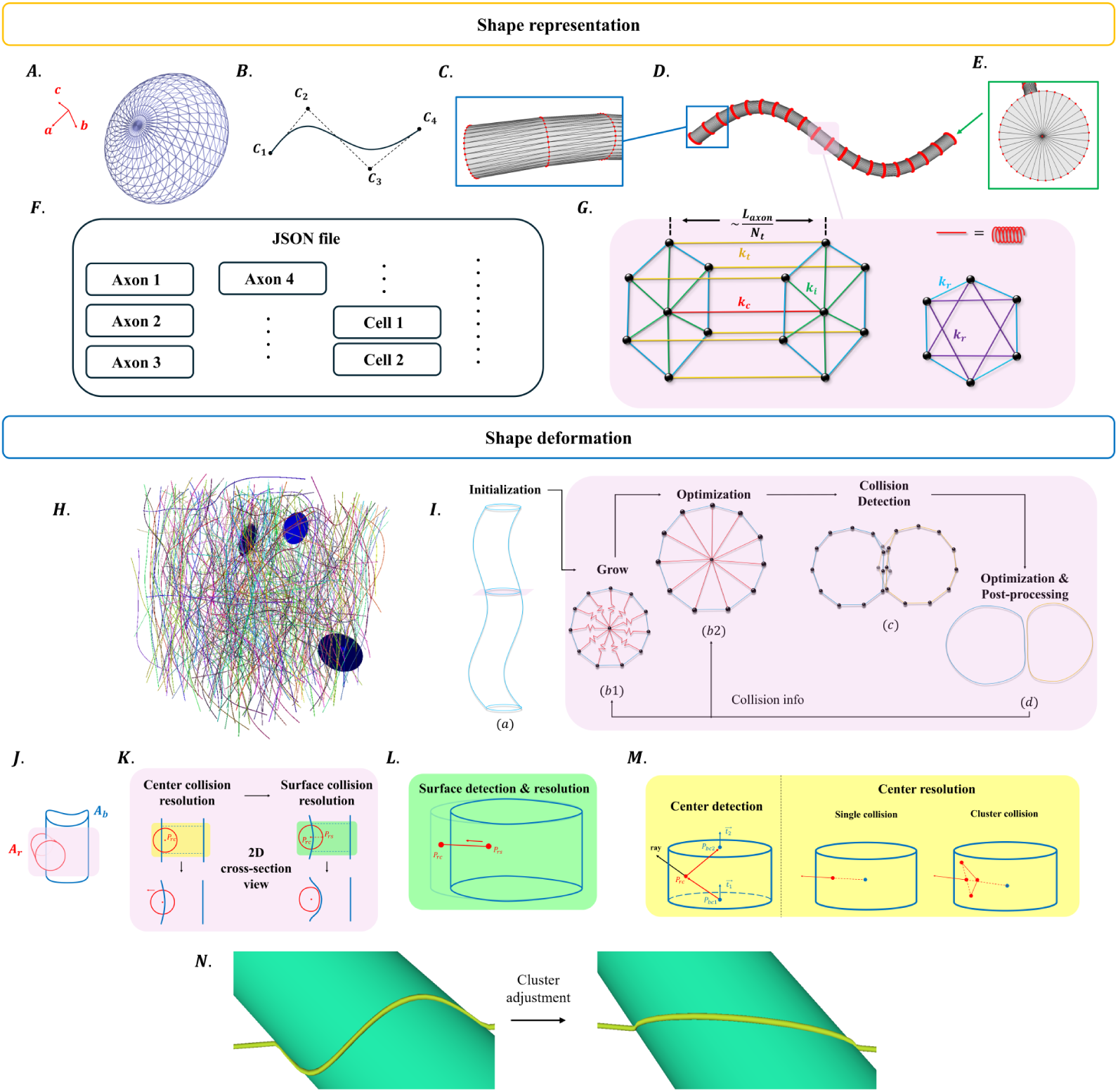
**A**. Mesh representation of a cell. **B**. An example of a NURBS curve with four control points. **C**. Zoomed-in view of the sampled point rings along the axon. A total of *N_t_* rings of points are sampled. **D**. Mesh representation of an axon. **E**. *N_r_*sampled points in the radial direction. **F**. Structure of the configuration JSON file. **G**. Mass-spring model defined using the sampled points as the mass points. Color-coded springs represent different mechanical components. **H**. An initialized voxel. **I**. Pipeline of the synthesis process. (*a*) Mesh representation and corresponding physical model initialized. (*b*1) The iterative growth (with pink background) begins with a growth step, during which vertex positions remain unchanged. (*b*2) Optimization step to realize growth and deformation. (*c*) Collision detection. (*d*) Another optimization step to update the shape according to the resolved collisions. **J**-**N**: Pipeline for collision detection. **J**. Schematic of the illustrated colliding scenario between axons *A_r_* and *A_b_*. **K**. 2D cross-section view of the expected solution for the center and surface point collision. **L**. Surface collision detection and resolution. **M**. Center collision detection and resolution. **N**. Effect of cluster adjustment of center collision resolution.

**Table 1:** Input parameters stored in the JSON configuration file for each cell body and axon. A complete voxel configuration contains one such entry for every structure in the synthesized substrate.

| **Cell** |  |  |  |
|  | **Name** | **Value type** | **Description** |
|---|---|---|---|
|  | *a* | Vector | Semi-major axis w/ length |
|  | *b* | Vector | Semi-intermediate axis w/ length |
|  | *c* | Vector | Semi-minor axis w/ length |
|  | *p<sub>c</sub>* | 3D Point | Coordinate of center point |
| **Axon** |  |  |  |
|  | **Name** | **Value type** | **Description** |
| **Structural Parameters** | control_points | Array of 3D points | Control points of the center curve of axons. A NURBS curve example illustrated in Figure 1.*B*. |
|  | *k* | Integer | Order of the center NURBS curve |
|  | weight | Array of Floats | Rational weight of NURBS curve associated with each control point |
|  | target_radii | Array of Floats | Maximum radii for axon growth associated with each control point |
|  | *g*-ratio | Array of Floats | *g*-ratio associated with each control point |
|  | *N<sub>myelin</sub>* | Integer | Number of myelin layers |
|  | *N<sub>t</sub>* | Integer | Number of sampled points in the axial direction |
|  | *N<sub>r</sub>* | Integer | Number of sampled points in the radial direction. The total number of sampled surface points of one axon is *N<sub>t</sub>* × *N<sub>r</sub>* |
| **Physical Parameters** | *κ<sub>r</sub>* | Array of floats / Float | Stiffness constant for springs on surface along the circumferential direction |
|  | *κ<sub>t</sub>* | Array of floats / Float | Stiffness constant for springs on surface along the axial direction |
|  | *κ<sub>i</sub>* | Array of floats / Float | Stiffness constant for springs connecting center to surface |
|  | *κ<sub>c</sub>* | Array of floats / Float | Stiffness constant for springs connecting center points |
| **Morphological Alteration Parameters** | *A<sub>beading</sub>* | Float | The amplitude of beading effect. Default is 0 (normal axon) |
|  | *f<sub>beading</sub>* | Float | The frequency of beading |
|  | is_loose | Boolean | Flag for myelin loosening |
| **Dynamic Parameter** | *N<sub>iter</sub>* | Integer | Number of iterations of growth. Larger number of iterations gives better quality while requires longer time to generate. Default is 10. |

Axon inputs comprise four groups: structural, physical, morphological alteration, and dynamic parameters. Structural parameters specify the centerline, target radius and *g*-ratio profiles, number of modeled myelin layers *N*_myelin_, and axial and circumferential sampling counts *N_t_*and *N_r_*. Physical parameters specify the stiffnesses of circumferential, axial surface, radial center-to-surface, and centerline springs. Morphological alteration parameters specify the beading amplitude *A*_beading_, frequency *f*_beading_, and the Boolean flag is_loose, which enables myelin layer decompaction. The single dynamic parameter *N*_iter_ controls the number of growth iterations, with a default of 10.

PhyMorphS constructs triangle meshes from the initial configuration. Cell bodies are represented as rigid ellipsoidal meshes, whereas axonal meshes deform during synthesis (Fig. 1.A and D). Each axon is initialized around a centerline described by a non-uniform rational B-spline (NURBS) curve (Piegl and Tiller, 2012). A NURBS curve is a parametric representation for modeling smooth trajectories (Piegl and Tiller, 2012). It is defined by a set of `control_points`, weight, a knot vector, and the order *k*. The control points determine the overall curve shape, the weights control the influence of individual control points, and the order *k* determines the smoothness of the curve, with a default value of 3. Fig. 1.B shows an example curve defined by four control points.

Each control point is also assigned a target radius and a *g*-ratio, which are stored in `target_radii` and *g*-`ratio`. These control-point values are evaluated along the centerline using the same NURBS settings to obtain smoothly varying target radius and *g*-ratio profiles. The target radii guide subsequent growth, while the final geometry also depends on mechanical and packing constraints.

To construct the mesh representation of the axon, we use user-defined *N_t_* and *N_r_* to control sampling along the axial and radial directions, respectively. We first sample *N_t_* points along the center curve to determine the center points. For each center point, we sample *N_r_* surface points uniformly on a circular cross-section with the initialized radius, defined in the plane orthogonal to the tangent direction of the curve. We denote the collection of the surface points on the same cross-section as a *ring*. In total, this results in *N_t_ N_r_*vertices for the surface mesh of a single axon. Together, these structural parameters fully define the mesh-based representation of the axonal geometry. Fig. 1.H shows an example of a voxel initialized from a given configuration.

### 2.3. Mass-spring system

We extend the mass-spring formulation introduced in (Cui et al., 2024) by revising spring connectivity and growth constraints to improve synthesis robustness under dense packing. The resulting system models axonal mechanics through springs representing membrane tension and internal structural support (Fig. 1.G). The mass-spring system is a model widely used in physics-based simulations to simulate deformable objects due to its simplicity, efficiency, and intuitive physical interpretation (Nealen et al., 2006). The shape is modeled using a collection of mass points connected by springs, which enables the simulation of elastic behavior under various external forces. Each spring follows Hooke’s law, stating that the force applied by the spring is proportional to its displacement from the equilibrium position.

Surface vertices and sampled center points serve as mass points, giving *N_t_*(*N_r_* + 1) mass points for an axon with *N_t_* rings and *N_r_* surface vertices per ring. Springs connect these points and generate forces proportional to their extension or compression relative to their rest lengths.

As shown in Fig. 1.G, within each ring, circumferential springs connect neighboring surface vertices (blue), while radial springs connect each surface vertex to the ring center (green). Additional springs connect surface vertices separated by one intervening vertex to resist bending (purple) (Baraff and Witkin, 2023). Between neighboring rings, axial surface springs connect corresponding surface vertices (yellow), and centerline springs connect the ring centers (red). The surface connections approximate membrane tension, while the radial and centerline connections provide effective internal support.

Each spring has a rest length *L*_0_ and a stiffness coefficient *κ*. Rest lengths are initialized from the Euclidean distances between connected mass points. The parameters *κ_r_*, *κ_t_*, *κ_i_*, and *κ_c_* control circumferential and bending, axial surface, radial, and centerline springs, respectively, as illustrated in Table 1. All springs use the same coefficient by default. Users can instead assign a scalar coefficient to each spring type within an axon or provide individual coefficients for its springs.

### 2.4. Iterative optimization

After initialization, axons are iteratively grown from their initial radii toward user-defined target radii. As axons expand, they may intersect, resulting in what we define as a *collision*. We therefore employ the iterative synthesis process illustrated in Fig. 1.I to detect and resolve collisions throughout growth.

The voxel is initialized as a collection of triangle meshes from the JSON configuration. Axons then grow incrementally toward their prescribed target radii over a user-defined number of iterations, *N*_iter_. Each iteration comprises a growth stage, optimization, collision detection and resolution, and a second optimization followed by geometric post-processing. Increasing *N*_iter_ produces smaller growth increments, allowing the geometry to adjust more gradually to mechanical and packing constraints. The whole synthesis terminates after the prescribed number of iterations.

Through repeated growth, collision detection, and constrained deformation, axons interact with their local environment and progressively develop their final cross-sectional shapes and trajectories, motivated by observations that axonal morphology is modulated by surrounding tissue structures (Andersson et al., 2020).

#### 2.4.1. Step I: axon growth

Similar to the radial growth approach used in CACTUS (Villarreal-Haro et al., 2023), this process begins with reduced axonal radii. In PhyMorphS, all axons are initialized with a common radius equal to one-fifth of the minimum target radius specified in the input configuration. However, growth in PhyMorphS is coupled with mass-spring based deformation, allowing axonal geometry to adapt to physical constraints and interactions with neighboring structures.

Growth is driven by increasing the rest lengths of the radial springs connecting ring centers to surface vertices. This approach avoids previously detected collisions, as no vertex motion occurs during the growth stage. Instead, we only modify the spring energy by increasing the rest lengths. The actual movement of vertices is handled in the subsequent optimization stage, where the colliding points’ motion is appropriately constrained. Meanwhile, it helps preserve axonal volume, as the resulting forces will push both the central mass point and the non-colliding surface away from regions of contact. The rest length of the springs connecting corresponding center points and surface points in the *i*th iteration is calculated by

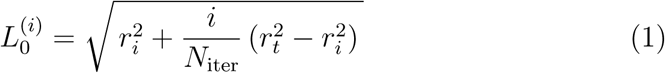

where *r_t_*is the target radius, *r_i_*is the initial radius, *N*_iter_ is the total number of iterations. This update performs a linear interpolation in *r*^2^, thereby ensuring volume-consistent growth across iterations.

In this setup, radial springs are set to a compressed state. During the subsequent optimization process, they progressively relax and expand, driving the growth of the corresponding axonal structures.

#### 2.4.2. Step II: optimization

Most applications of the mass-spring system focus on real-time dynamic simulation, in which velocity is explicitly modeled. In contrast, neural growth is a long-term process that can be approximated as a sequence of quasi-static equilibrium states. Therefore, we iteratively solve for the equilibrium configuration corresponding to the minimum system energy, without explicitly introducing velocity terms. According to the principle of minimum energy (Landau et al., 2012; Baraff and Witkin, 2023), within this conservative system, we determine the equilibrium by minimizing the total potential energy *E_s_*of the springs, i.e.

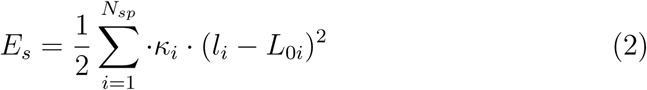

where *N_sp_*is the number of springs, *κ_i_, l_i_* and *L*_0*i*_ are the spring constant, current length and the rest length of the *i*th spring, respectively.

The parameters to be optimized are the coordinates of each sampled mass point. Let *S_m_* denote the set of springs incident to mass point *p_m_*, and let *n*(*i, m*) denote the index of the opposite endpoint of spring *i*. Their gradients can be easily obtained by the following equation:

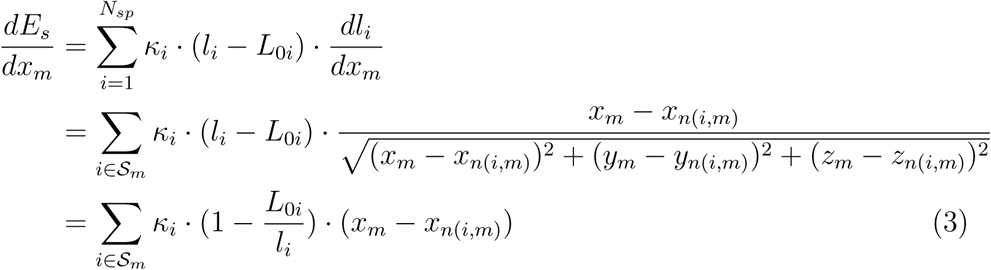

where *x_m_* is the *x* coordinate of mass points indexed *m*. The same calculation applies to the *y* and *z* coordinates.

However, several additional constraints on the updates of the mass point coordinates are needed to achieve greater realism. First, we restrict the motion of mass points involved in collisions detected in the previous iteration. Specifically, those colliding mass points are only allowed to move towards the center mass point of the ring, i.e.

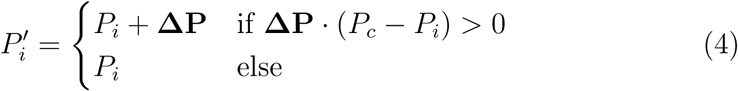

where *P_i_* denotes the coordinate of the mass point, *P_c_* is the coordinate of the central mass point in the same ring, and **ΔP** is the offset solved by optimization. This prevents the model from reverting to an equilibrium state that would otherwise cancel out the effects of collision resolution.

Furthermore, we project the gradients of each mass point onto the transversal plane (i.e., the plane perpendicular to the local tangent). Since axons grow only in the radial direction after forming connections, movement along the axial direction is undesirable. On the other hand, it also enhances the robustness of the model by preserving the ring order and preventing unintended axial shifts.

Another challenge is controlling spring length to avoid springs that are too long or too short. We define a barrier energy as follows

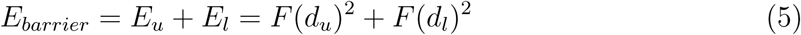

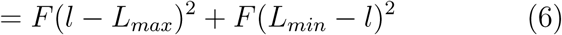

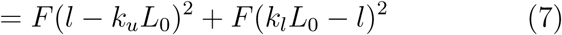

where *E_u_* and *E_l_* denote the energy for the upper barrier and the lower barrier. *F* (*x*) is an energy function defined to punish spring distance exceeding either barrier. *d_u_* and *d_l_* denote the differences between the current spring length *l* and the upper and lower barrier thresholds, respectively. *L_max_* and *L_min_* define the upper and lower barrier lengths, which are computed as fixed ratios of the spring rest length *L*_0_, where *k_u_* and *k_l_* are user-defined constants. In our experiments, we set *k_u_* = 2.0 and *k_l_* = 0.2

The energy function *F* (*x*) is defined as

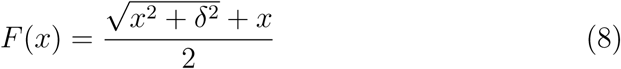

The smoothing parameter *δ* has units of length. The function *F* smoothly approximates max(*x,* 0), so the penalty is small inside the prescribed interval and increases approximately quadratically with violations beyond the thresholds. It therefore discourages excessive deformation without imposing hard length bounds. We set *δ* = 10*^−^*^6^ in the experiments.

The gradient is derived as follows

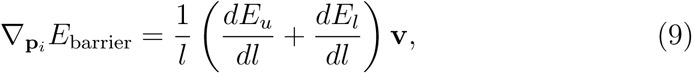

where,

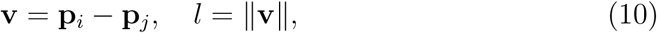

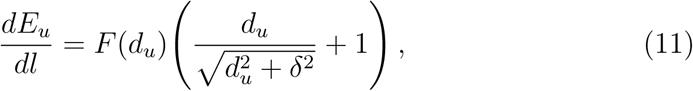

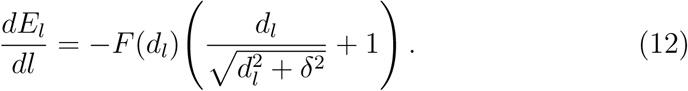

Thus, the optimization objective at this stage is given by the energy

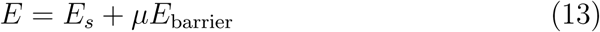

where *µ* denotes the weight of the barrier energy. In our experiments, we empirically set *µ* = 5 10^3^.

We employ the L-BFGS algorithm (Liu and Nocedal, 1989) as the solver. Although the system Hessian is sparse, explicitly forming or storing its inverse would generally lead to a dense representation and prohibitive memory consumption. L-BFGS avoids this issue by implicitly approximating the inverse Hessian in a limited-memory manner, making it well-suited for large-scale problems while maintaining fast convergence.

#### 2.4.3. Step III: collision detection

After growth, new collisions might have been introduced into the generated voxel. Thus, collision detection is required in each iteration to remove new collisions, as intersection-freeness is one property required on the input mesh for the downstream dMRI simulation.

In this work, we use the so-called broad phase/narrow phase approach for collision detection (Ericson, 2004).

##### Broad phase

In the broad phase, it uses simplified representations, such as bounding boxes, spheres, or spatial partitioning structures, to rapidly identify a subset of potential colliding candidates. This operation significantly reduces the number of geometric checks required in the subsequent narrow phase. In this work, we compute the axis-aligned bounding boxes (AABB) of cells and the closed shape formed by the neighboring rings (as shown on the left side in Fig. 1.G). Notably, we collect all the AABB pairs that are not adjacent in the same axon. In other words, we account for self-intersections, excluding only those pairs where both AABBs contain the same ring.

##### Narrow phase

After identifying the colliding AABB pairs in the previous stage, a finer per-point collision resolution is performed to handle all possible intersections. Fig. 1.J-N show a detailed illustration of the narrow phase. We set a simple scenario: A red axon *A_r_* collides with a blue axon *A_b_*. Fig. 1.K shows a simple two-step pipeline for one AABB pair.

Firstly, a center collision resolution is conducted. The detection is done using the typical ray casting algorithm (De Berg, 2000). We shoot a ray with random direction from *P_rc_* and count the number of ray-triangle intersections. If the number of intersections is odd, *P_rc_* lies within this volume. Otherwise, it is outside. After detection, the collision resolution is not immediately performed. We collect all the collision information for all rings in one axon and then handle it by dividing into two cases. If only one isolated center point of a ring is involved in a collision (meaning that no center points of its neighboring rings collide), *P_rc_* is displaced away from the corresponding blue center point as shown in Fig. 1.M. Otherwise, we consider all neighboring colliding center points as a cluster. An auxiliary line is drawn between the first point and the last point in the ordered cluster. The displacement direction of the cluster is along the perpendicular bisector of the auxiliary line that passes through the blue center point. In addition, this blue center point in Fig. 1.M is at the average position of all corresponding center points of the cluster. With the displacement direction determined, we move *P_rc_* accordingly along that direction. And this is referred to as *cluster adjustment*. Fig. 1.N illustrates its effect.

One reason for solving center collisions first is that these collision-free centers can determine the moving direction of the surface points. In the surface collision resolution stage, instead of calculating the displacement direction, we shrink the colliding region towards the center. Thus, for each surface point, we detect the collision by solving segment-triangle intersections, i.e., segment 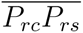 and the triangle mesh of the volume surface as shown in Fig. 1.L. If it has intersections, this surface point should be displaced towards the center.

For axon-axon collisions, the response is symmetric, with both axons displaced to resolve collision, as indicated in Fig. 1.K. Because each displacement changes the contact geometry, collision detection and displacement are repeated iteratively until the detected collisions are resolved.

For axon-cell collisions, cells are treated as fixed ellipsoidal solids. We test whether axonal center and surface points lie inside the ellipsoid and resolve collisions by displacing only the affected axonal points. In both cases, displacements of center and surface points are projected onto their respective transversal planes.

The collision framework can also be extended to other structures represented by closed triangle meshes, with displacement rules adapted to their geometry and motion constraints.

#### 2.4.4. Step IV: optimization and post-processing

Because collision resolution can perturb the mechanical equilibrium, we perform an additional potential-energy minimization step to restore physical realism. Meanwhile, some post-processing procedures are performed to enhance the robustness of the iterative pipeline. We first check if all the mass points still lie on their own transversal planes; if not, we project them. Then we perform smoothing on the surface points to avoid artificial sharp corners.

### 2.5. Morphological alteration

#### 2.5.1. Axonal swelling

Axonal swelling is characterized by an increase in axonal diameter, producing enlarged regions that may be localized or extend over longer segments. These regions vary in their degree of enlargement, longitudinal extent, and cross-sectional shape. In PhyMorphS, target_radii is an array of floatingpoint values in the configuration file, with one entry specifying the local target radius associated with each centerline control point as shown in Table 1. Users prescribe swelling by increasing selected entries in this array.

#### 2.5.2. Axonal beading (pearling)

Axonal beading is characterized by repeated swellings separated by constricted segments along an axon. We model this pattern by sinusoidally modulating the rest lengths *L*_0_ of radial springs connecting ring centers to surface points. The user-defined parameters *A*_beading_ and *f*_beading_ control the modulation amplitude and frequency, respectively. Both parameters are specified per axon and are zero by default, allowing users to introduce beading selectively. Subsequent optimization and collision resolution determine the resulting beaded geometry under local packing constraints.

#### 2.5.3. Myelin layer decompaction

Myelin is modeled as multiple concentric mesh layers tightly wrapped around the axon, following the multi-layer representation proposed by Canales-Rodríguez et al. (2024). Users can customize the initial configuration to specify the *g*-ratio, defined as the ratio of the inner axonal diameter to the outer fiber diameter, and the number of myelin lamellae, *N_myelin_*. In normal configuration, the myelin layers are tightly spaced to form a compact sheath. We additionally support the synthesis of myelin decompaction (Corral-Bolaños et al., 2026), in which the inner layers exhibit increased interlamellar spacing. To introduce morphological variability, each myelin layer can also be assigned a random cross-sectional morphology.

### 2.6. Implementation details

#### 2.6.1. Preliminary centerline optimization

Before constructing the mesh-based mass-spring model, we perform a preliminary centerline optimization to reduce initial overlaps and accelerate sub-sequent deformation during growth. Inspired by the existing primitive-based methods (Villarreal-Haro et al., 2023; Winther et al., 2024b; Ginsburger et al., 2019; Callaghan et al., 2020; Close et al., 2009), we represent each axon as a chain of spheres defined by their centerline positions and corresponding target radii. The neighboring spheres along each axon are connected by virtual springs. This initialization is performed once to directly modify the configuration file, before it is fed into PhyMorphS framework.

We implement the preliminary optimization using an approach analogous to the mass-spring system. The center positions of spheres are optimized using Adam to minimize an energy comprising four terms: a collision term penalizing sphere overlaps, a boundary term penalizing departures from the voxel domain, a connectivity term penalizing deviations from the rest lengths between neighboring spheres, and a bending term penalizing curvature to promote smooth centerlines. The endpoints on the voxel boundaries remain fixed throughout this preliminary optimization. At each iteration, the gradients are computed through automatic differentiation and projected to satisfy the boundary constraints before updating the sphere centers. The target radii guide this initialization. Subsequent deformation in PhyMorphS under mechanical and packing constraints can produce final radii that differ from these targets.

#### 2.6.2. Parallelization and collision resolution

The optimization in the iterative pipeline is performed separately for each axonal mass-spring system, enabling parallel computation. Separate solves allow other axons to continue optimizing when an individual solve stalls in a local minimum or terminates early, improving robustness compared with whole-scene optimization. Collision computations are also parallelized over candidate AABB pairs.

During all but the final growth iteration, conservative displacement updates replace full iterative collision resolution, reducing repeated collision checks while providing an approximate response during intermediate growth. Full iterative collision resolution is performed only once in the final growth iteration. The whole process terminates after the user-defined *N*_iter_ iterations.

#### 2.6.3. Additional morphological variability

During growth, axonal deformation arises from axon-axon interactions, axon-cell interactions, and mass-spring optimization. However, rings away from regions of frequent contact tend to retain circular cross-sections. Interactions with blood vessels and long-term deformations caused by cell movement are also not explicitly modeled. To account for additional morphological variability, we apply Perlin noise (Perlin, 1985), a procedural noise function that generates natural-looking surface variations, after the growth loop. A global noise amplitude of 0.5, selected empirically, is used consistently across all experiments.

### 2.7. Experiment settings

#### 2.7.1. Real data and synthesis settings

We evaluated PhyMorphS using segmentations of 3D XNH images of vervet monkey brain white matter (Amirrashedi et al., 2025). The segmented data comprise thousands of 3D axons from four WM regions: crossing fibers (CF) in the centrum semiovale, the splenium and midbody of the corpus callosum, and U-fibers beneath the cortex. The data were acquired at 75-100 nm resolution with fields of view extended up to 200 200 700 µm^3^, capturing axons up to 700 µm in length. Fiber orientation distributions (FODs), axon diameter distributions (ADDs), and cross-sectional morphological statistics derived from these data provide the basis for both initialization and evaluation.

For each region, we generate 10 cubic voxels with side lengths of 50 µm. We set the target FVF to 80% as a deliberately high packing target to evaluate the ability of PhyMorphS to synthesize densely packed substrates. The target FVF guides initialization and is not necessarily reached after synthesis. Initial configurations are sampled from the fitted regional FODs and ADDs. After iterative optimization, the synthesized morphology is quantitatively compared with the real data.

To demonstrate morphological alterations, we assign axonal beading or myelin layer decompaction to selected axons and qualitatively assess the resulting shapes. The reference tissue consists of 1-mm biopsies from the corpus callosum of mice, stained with uranyl acetate and lead citrate and examined using a CM100 TEM microscope (Corral-Bolaños et al., 2026).

#### 2.7.2. Fit orientation and diameter distributions

To initialize the configuration file using the real data, we need to fit the distributions for both fiber orientation and axon diameter, and then draw samples from the fitted distributions.

##### Fiber orientation distribution (FOD)

For each WM region, we estimate one regional FOD from the measured axonal orientations. This distribution defines the probability of sampling a given axonal orientation during voxel initialization. It does not prescribe the spatial organization of fibers within the voxel. We estimate each regional FOD from the measured orientations using an antipodally symmetric von Mises-Fisher kernel density estimator with concentration parameter of 100 (Banerjee et al., 2005). This parameter controls angular smoothing, with larger values producing narrower kernels. The estimate is evaluated on a dense spherical grid and normalized by its maximum before fitting a real spherical harmonic (SH) expansion with maximum degree of 28 (Axler et al., 2013). Only even degrees are included, preserving antipodal symmetry and treating opposite directions as equivalent axonal orientations. This representation accommodates multiple preferred fiber orientations within each region.

##### Axon diameter distribution (ADD)

Imaging and segmentation limitations cause thin axons to be underrepresented in the XNH measurements (Amirrashedi et al., 2025). Thus, we treat the measured XNH diameters as a left-truncated observation of the full ADD distribution and estimate the full ADD using a gamma distribution *f* (*d|α, θ*).

Because segmentation provides no explicit detection threshold, we first fit an auxiliary shifted gamma distribution *f_t_*(*d|α_t_, θ_t_,* loc) to the observed diameters and use its mode as an empirical cutoff. For *α_t_ >* 1, this gives

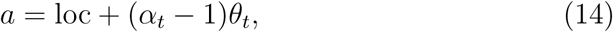

where *α_t_*, *θ_t_*, and loc are the fitted shape, scale, and location parameters, respectively.

We then retain only the *n* measurements satisfying *d_i_ > a*. Their truncated likelihood is

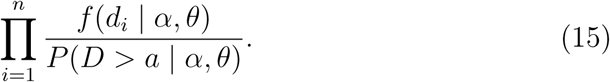

The underlying gamma parameters are estimated by minimizing the corresponding negative log-likelihood:

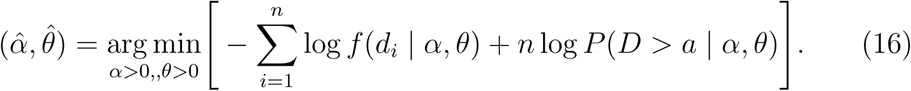

The resulting full gamma distribution is used for initialization. Its lower tail below *a* is extrapolated from the fitted model.

#### 2.7.3. Sampling and voxel initialization

Following a strategy similar to the two-dimensional packing approach of Hall and Alexander (2009), we first estimate the number of axons required for the target FVF using the fitted regional ADD. We then generate their initial parameters through Monte Carlo sampling. For each axon, we draw an orientation and a diameter *d* from the corresponding fitted regional distributions, FOD and ADD, and assign a target radius *r* = *d/*2. A random point within the voxel determines the axon’s spatial position, and control points are sampled along the assigned orientation to define its centerline. The resulting parameters are stored in the JSON configuration file.

### 2.8. Evaluation metrics

We assess the final synthesized substrates in terms of packing density, axonal size, longitudinal diameter variation, cross-sectional shape, and tortuosity.

#### Fiber volume fraction (FVF)

We quantify fiber packing as FVF = *V*_fiber_*/V*_voxel_, where *V*_fiber_ is the total volume occupied by axons and their myelin sheaths within the voxel, excluding cell bodies, and *V*_voxel_ is the total voxel volume. For each closed axonal or myelin mesh, the enclosed volume is computed analytically from its triangle mesh representation, and the volumes of all fibers are summed to obtain *V*_fiber_. Higher FVF values indicate denser fiber packing.

#### Axon diameter distribution (ADD)

For each cross-sectional ring sampled along an axon, we calculate the equivalent circular diameter as 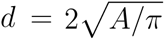, where *A* is the area enclosed by the ring (Abdollahzadeh et al., 2019). The transversal motion constraints keep the vertices of each ring coplanar. We calculate the mean diameter of the rings for each axon and pool the mean diameters to construct ADD histograms and fit gamma distributions. Full ADD distributions are compared between initialized and synthesized data. For comparison with the XNH measurements, we additionally apply the truncation threshold *a* to remove axons whose mean diameter is smaller than *a* and compare the truncated histograms.

#### Per-axon coefficient of variation of diameter

For each axon, we calculate 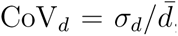, where *σ_d_* and 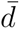 are the standard deviation and mean of its ring diameters, respectively. This dimensionless measure quantifies longitudinal diameter variation relative to the mean diameter. A constant diameter gives CoV*_d_* = 0, while larger values indicate greater relative variation. The resulting histogram contains one value per axon.

#### Cross-sectional eccentricity

Following Winther et al. (2024b), we apply principal component analysis (PCA) to each ring’s vertices and obtain the two in-plane covariance eigenvalues, *⋋*_1_ *≥ ⋋*_2_ > 0. The effective semimajor and semiminor axis lengths are proportional to 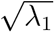 and 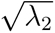, respectively, giving the eccentricity 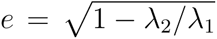. Values near zero indicate similar spread along the two principal directions, whereas values approaching one indicate strongly elongated cross-sections.

#### Cross-sectional convexity

We quantify cross-sectional convexity as *C* = *A/A_h_*, where *A* is the area enclosed by the ring and *A_h_*is the area of its convex hull. Values satisfy 0 *< C* 1. Convex cross-sections have *C* = 1, while lower values indicate a larger relative area deficit within the convex hull and therefore greater concavity.

#### Tortuosity

Tortuosity is defined as *T* = *L/L_s_*, where *L* is the axonal centerline length, approximated by summing segment lengths along the spline, and *L_s_* is the straight-line distance between its endpoints. A straight trajectory has *T* = 1, while larger values indicate greater path length relative to endpoint separation.

## 3. Results

### 3.1. Regional substrate synthesis and packing density

We synthesized 10 (50 µm)^3^ voxels for each WM region using the corresponding ADD and FOD estimated from the real XNH data. The synthesized substrates are densely packed within the voxel boundaries (Fig. 2.A). Consistent with the regional FODs, axons in the splenium and midbody are predominantly parallel with few crossings, whereas the CF region contains clearly intersecting fiber populations and the U-fibers exhibit a broader range of orientations. Across all regions, the visual diameter distribution is dominated by smaller axons with fewer large axons, consistent with the fitted regional ADDs.

**Figure 2:**
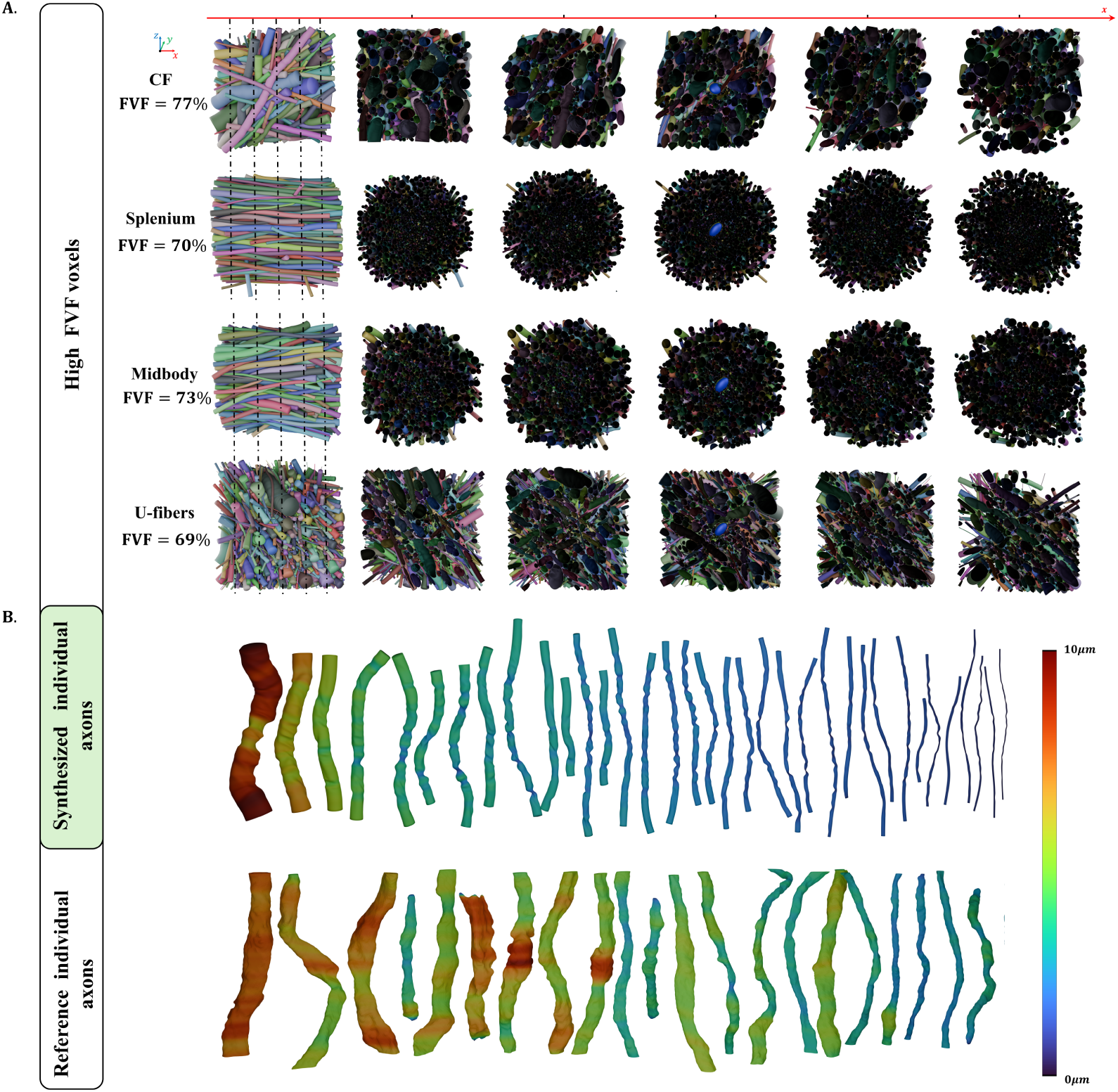
**A.** Visual results of the synthesized voxels and their cross-sectional view along the x-axis. The leftmost panel of each row shows the fiber-bundle arrangement. The remaining panels show cross-sections sampled at successive positions along the x-axis at the marked positions. A fixed cell (blue ellipsoid) is placed in the center of the voxel. The target FVF is set to 80%. **B.** Individual axons comparison between synthesized results (top) and segmentation data (bottom).

With a target FVF of 80%, the achieved FVF across the 10 realizations was 79.14 ± 2.26% for CF, 72.24 ± 0.92% for the splenium, 73.85 ± 1.61% for the midbody, and 65.82 ± 4.15% for U-fibers. Thus, the same target FVF results in region-dependent final packing densities.

Visual inspection of selected synthesized axons and morphologically comparable XNH-segmented axons from the CF region shows similar ranges of diameter, longitudinal caliber variation, and trajectory curvature (Fig. 2.B). In both datasets, larger axons exhibit more pronounced absolute diameter variation along their trajectories, whereas smaller axons show less variation. Additionally, the synthesized set spans smaller diameters because it represents the estimated full ADD, whereas small axons are underrepresented in the XNH segmentation (Amirrashedi et al., 2025).

### 3.2. Morphological agreement with the real data

We compare the synthesized and XNH morphologies using ADD, CoV*_d_*, convexity, eccentricity, and tortuosity. Distributional differences are quantified using the Wasserstein distance (Villani et al., 2009). Smaller Wasserstein distances indicate greater similarity between the two distributions.

#### 3.2.1. Axon diameter distribution

The estimated full ADD extends below the region-specific truncation threshold *a*, accounting for the small-diameter population that is incompletely observed in the segmentation (Fig. 3.A, left column).

**Figure 3:**
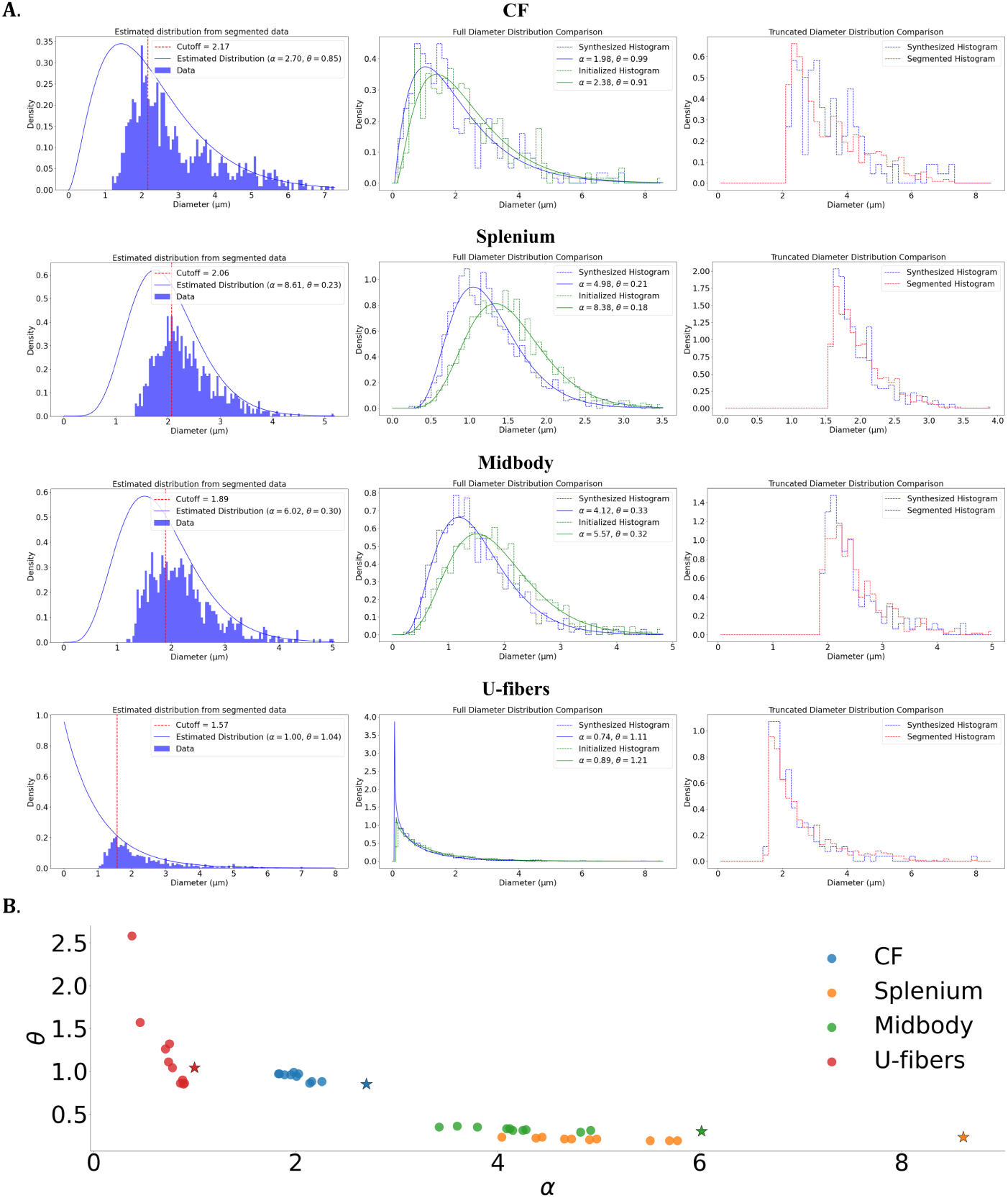
**A.** Axonal diameter distribution across different WM regions for initialized, synthesized, and segmented (real) data. **Left column**: the estimated distribution from the truncated real data. **Middle column**: The full distribution functions of initialized (green) and synthesized (blue) data. **Right column**: The truncated data histogram of synthesized (blue) and real (red) data. **B.** Scatter plot of the (*α, θ*) parameters for estimated ADD from real data (stars) and ADD of the synthesized data (circles) across four different regions.

Relative to the initialized ADD, the synthesized ADD shifts toward smaller diameters in all four regions (Fig. 3.A, middle column). The fitted mean diameter, *αθ*, decreases from 2.16 to 1.96 µm for CF, from 1.51 to 1.04 µm for the splenium, from 1.78 to 1.36 µm for the midbody, and from 1.08 to 0.82 µm for U-fibers, corresponding to reductions of approximately 10–31%.

Applying the same region-specific truncation thresholds to the synthesized data yields diameter distributions comparable to those observed in the real XNH segmentation over the reliably segmented diameter range (Fig. 3.A, right column). The fitted gamma parameters of the synthesized voxels are systematically shifted relative to the estimated reference ADDs, particularly in *α* (Fig. 3.B).

#### 3.2.2. Longitudinal diameter variation

Longitudinal diameter variation is quantified using the per-axon coefficient of variation, CoV*_d_* (Fig. 4.A). The distributions are predominantly below 0.5 in all four regions. The Wasserstein distances between synthesized and XNH distributions are 0.0500 *±* 0.0054 for CF, 0.0420 *±* 0.0041 for the splenium, 0.0387 *±* 0.0029 for the midbody, and 0.0333 *±* 0.0030 for U-fibers.

**Figure 4:**
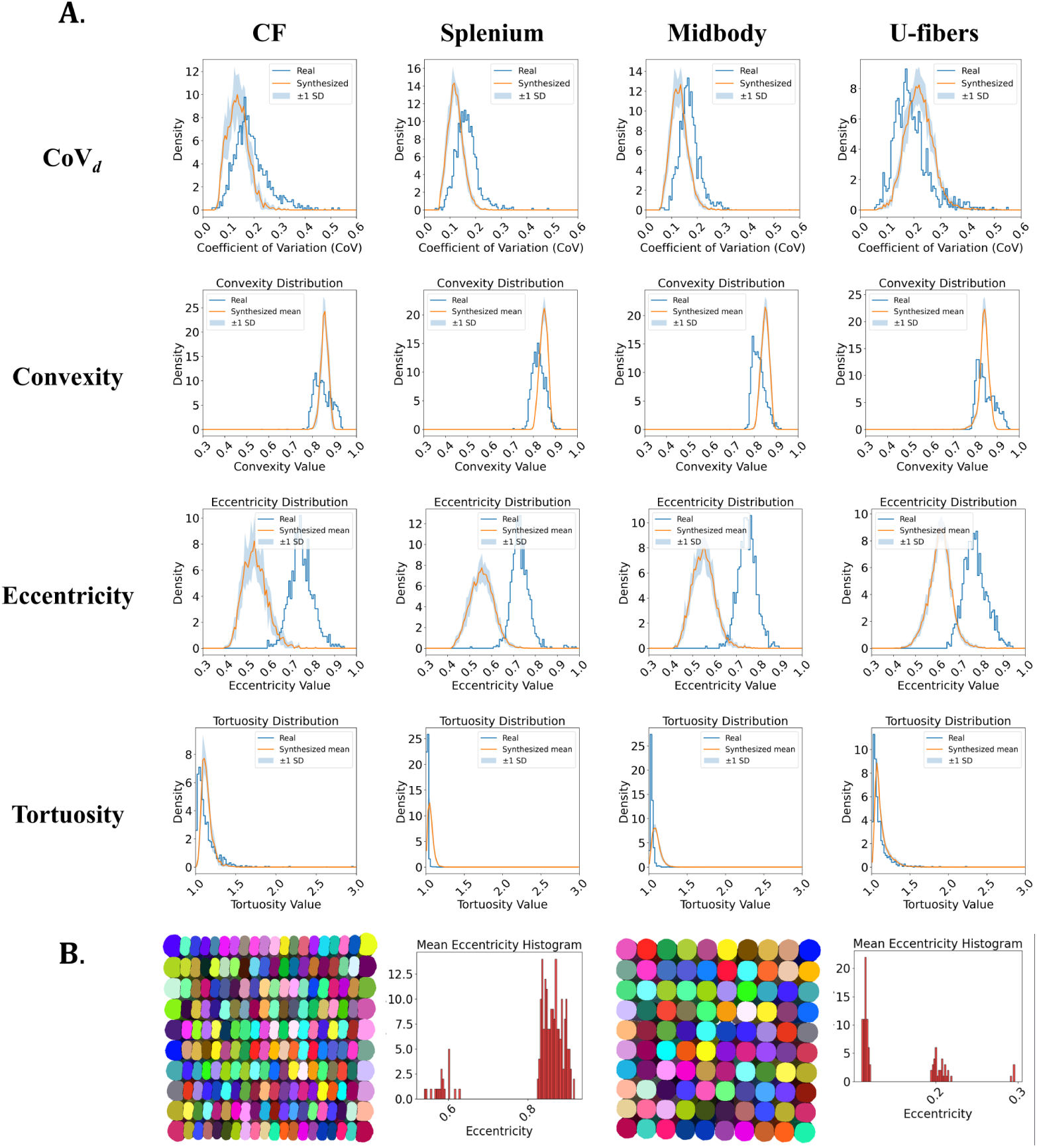
**A.** Distributions of CoV*_d_*, convexity, eccentricity, and tortuosity across four WM regions (CF, splenium, midbody, and U-fibers). For the synthesized data (orange), solid curves show the mean distribution across 10 independently generated voxels, and shaded regions indicate ±1 standard deviation (SD) across voxels. Real data from XNH are shown in blue. **B.** Example illustrating the influence of bundle organization on eccentricity. The figure and plot on the left show an anisotropically packed bundle and its corresponding eccentricity. The bundle is initialized with 20 axons along the x-direction and 10 axons along the y-direction. The figure and plot on the right show a uniformly initialized scene, resulting in a lower eccentricity.

#### 3.2.3. Cross-sectional morphology

Convexity shows relatively small differences between synthesized and XNH distributions (Fig. 4.A), with Wasserstein distances of 0.0207 ± 0.0012 for CF, 0.0221 *±* 0.0009 for the splenium, 0.0296 *±* 0.0025 for the midbody, and 0.0176 *±* 0.0005 for U-fibers.

Eccentricity exhibits a larger discrepancy, with the synthesized distributions shifted toward lower values relative to the real XNH data (Fig. 4.A). The corresponding Wasserstein distances are 0.2106 *±* 0.0051 for CF, 0.1748 *±* 0.0058 for the splenium, 0.1996 *±* 0.0035 for the midbody, and 0.1601 *±* 0.0022 for U-fibers.

To examine the effect of fiber arrangement on cross-sectional eccentricity, we compared two controlled packing configurations (Fig. 4.B). An anisotropic arrangement with 20 axons along one direction and 10 along the orthogonal direction produces higher eccentricity than a uniformly initialized arrangement. This demonstrates that cross-sectional eccentricity depends on the initialized fiber arrangement, in addition to the interaction-driven deformation introduced by the PhyMorphS framework.

#### 3.2.4. Tortuosity

The XNH tortuosity distributions are concentrated near 1.0 in the splenium and midbody and are broader in CF and U-fibers (Fig. 4.A). The synthesized data reproduce the same regional trend, although they tend to exhibit slightly higher tortuosity in the relatively straight splenium and midbody regions and lower tortuosity in the more curved CF and U-fiber regions. The Wasserstein distances are 0.0448 *±* 0.0050 for CF, 0.0416 *±* 0.0073 for the splenium, 0.0630 *±* 0.0099 for the midbody, and 0.0306 *±* 0.0064 for U-fibers.

### 3.3. Morphological alterations

PhyMorphS supports user-defined morphological alterations, demonstrated here using axonal beading and myelin layer decompaction.

#### 3.3.1. Axonal beading

Fig. 5.A shows four axons synthesized within the same voxel with different combinations of *A*_beading_ and *f*_beading_: (0.8, 10), (0.2, 1.0), (0.4, 2.5), and (0.6, 5), respectively. Interactions with neighboring structures constrain the prescribed swellings, producing local variations in their size and shape. To isolate the effect of each parameter, we additionally synthesize single axons without neighboring structures (Fig. 5.B). Holding *A*_beading_ fixed while decreasing *f*_beading_ from 10 to 1 increases the spacing between swellings, whereas holding *f*_beading_ fixed while decreasing *A*_beading_ 0.8 to 0.2 reduces their magnitude.

**Figure 5:**
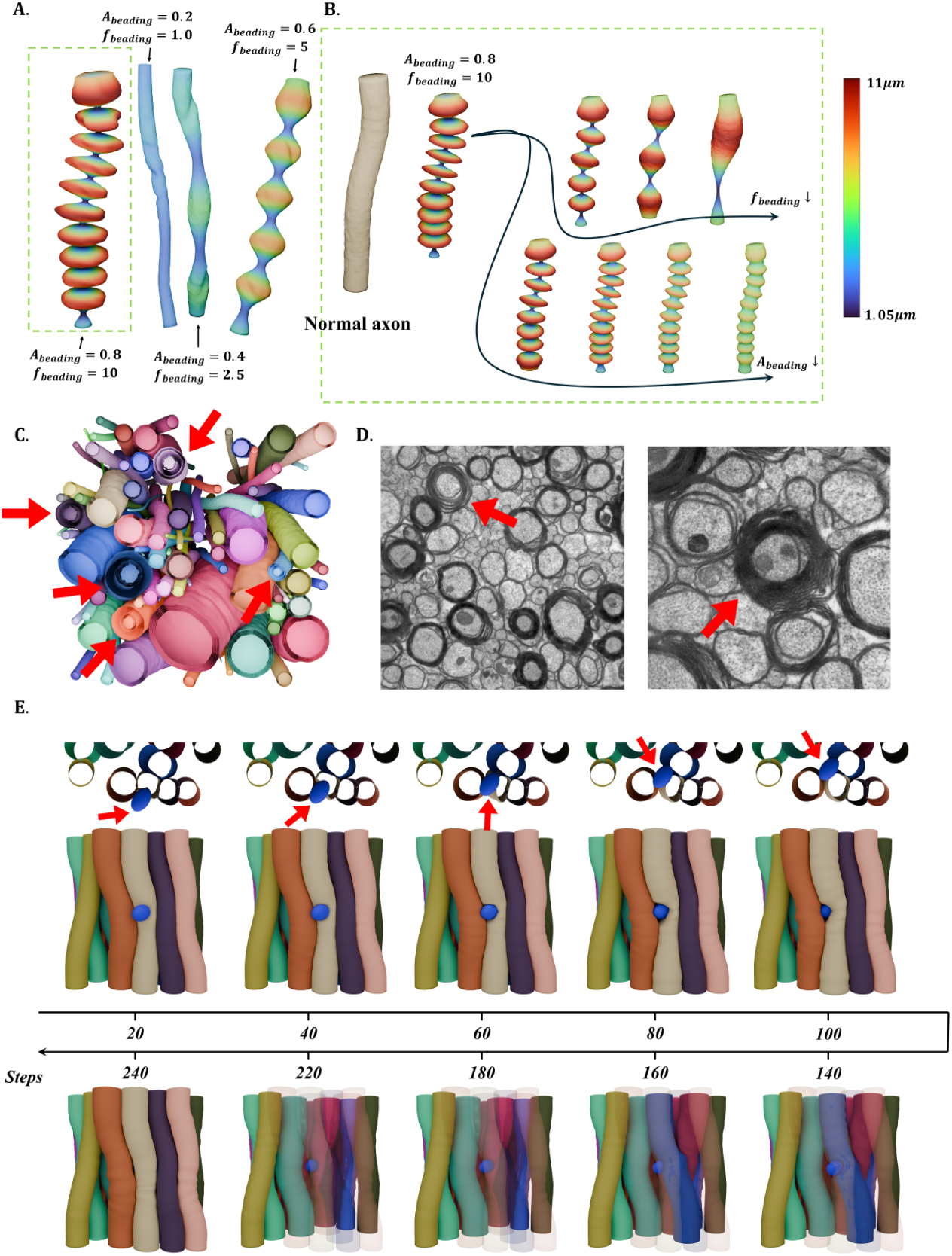
**A.** Four different axons in one scene with different *A_beading_*and *f_beading_*. **B.** A parameter study of beading parameters. The axon on the left is the normal axon without the beading effect. It corresponds to the beading axon on the left. The top row visualizes shape changes while decreasing *f_beading_*. The bottom row shows the changes while decreasing *A_beading_*. The axons are colored by their local diameters. **C.** A voxel with myelinated axons. Among them, five indicated axons are modeled with myelin layer decompaction (red arrows). **D.** Two representative EM images showing axons with loosened myelin, indicated by red arrows. **E.** Post-growth dynamic system. A moving cell (blue ellipsoid) traverses the scene along the x-axis, inducing additional deformation of neighboring axons through interaction. The top row shows a top-view cross-section of the affected region in the first 100 iterations, with the cell indicated by red arrows. Axons previously contacted by the moving cell are rendered transparent to visualize the deformation path.

#### 3.3.2. Myelin layer decompaction

Myelin layer decompaction produces increased spacing between inner myelin layers and a less compact sheath (Fig. 5.C). In this demonstration, each axon is represented using *N*_myelin_ = 5 layers. Myelin layer decompaction is enabled for five selected axons by setting is_loose=true, while compact myelinated axons use is_loose=false. The synthesized decompacted sheaths exhibit visibly increased and irregular spacing between the inner myelin layers. A similar separation of adjacent myelin lamellae is visible in the representative real EM images in Fig. 5.D, particularly in the regions indicated by the red arrows (Corral-Bolaños et al., 2026).

### 3.4. Post-growth dynamic deformation

Post-growth cell movement produces a spatially propagating deformation of the surrounding axons (Fig. 5.E). As the cell traverses the bundle, axons in direct contact are displaced and locally bent around the cell, while neighboring axons undergo smaller displacements through mechanical coupling. After the cell has passed, the directly affected axons partially return toward their pre-contact positions. In this experiment, the cell moves from 50 µm to 50 µm over 400 time steps. Axons in direct contact deform during passage and partially recover after the cell moves away.

### 3.5. Computational efficiency

Synthesis time scales approximately linearly with the number of axons over the evaluated range (Fig. 6), demonstrating favorable scalability of the parallelized optimization and collision-resolution pipeline. The largest tested substrate, (150 µm)^3^ with more than 2000 axons, is synthesized in approximately 3 hours. The efficiency experiments use CF configurations, whose heterogeneous orientations produce frequent collision checks, with voxel side lengths ranging from 25 µm to 150 µm and a target FVF of 80%. All experiments are run using 20 CPU threads on an Intel Xeon E5-2650 v4 processor (12 cores/24 threads, 2.2 GHz).

**Figure 6:**
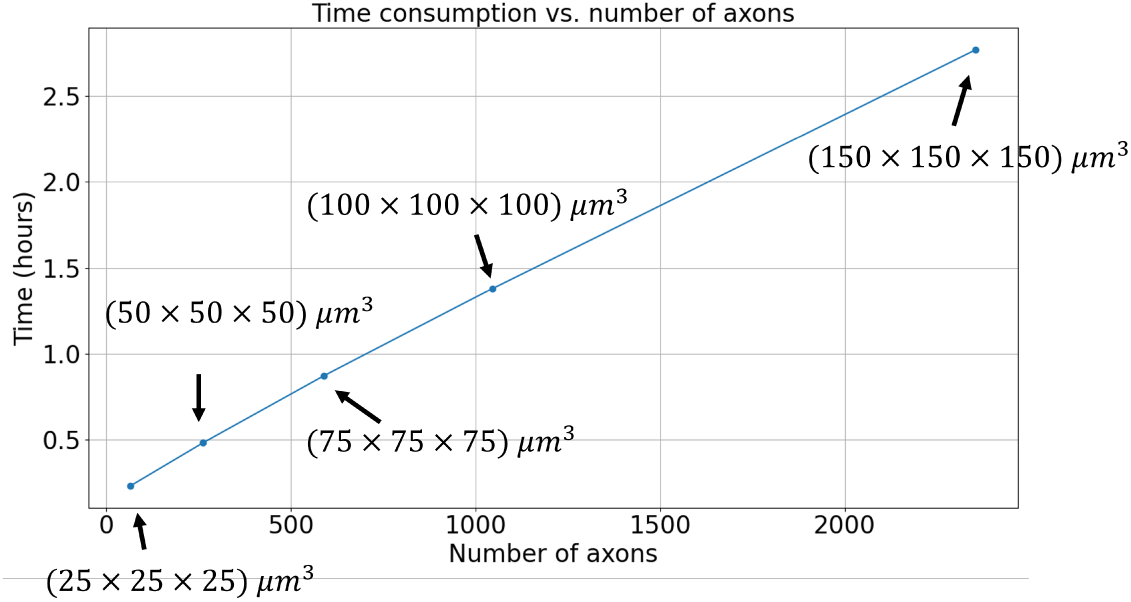
Computational efficiency plot using voxels generated from data of CF.

## 4. Discussion

We present PhyMorphS, a physics-guided, mesh-based framework for synthesizing WM microstructure, motivated by the observation that axonal morphology is modulated by interactions with the local tissue environment Andersson et al. (2020). Existing synthesizers often initialize axons from predefined geometric primitives and subsequently modify their shapes through growth and packing procedures. In contrast, PhyMorphS uses a mass–spring system to drive the mechanical deformation of triangle meshes, allowing axons to develop asymmetric, non-convex, and spatially varying morphologies under explicit mechanical constraints, consistent with those observed in 3D XNH data Amirrashedi et al. (2025); Andersson et al. (2020). Evaluation across four WM regions shows that the synthesized axons reproduce regional characteristics in diameter variation, cross-sectional convexity, and tortuosity, while also demonstrating the influence of tissue organization on eccentricity and achievable packing density. PhyMorphS further supports user-defined morphological alterations, including axonal beading and myelin layer decompaction, while preserving interactions with surrounding structures. Its explicit closed-mesh substrates provide known geometric ground truth for diffusion simulations, evaluation of biophysical models, and generation of paired microstructure-signal datasets for training machine-learning models.

### 4.1. PhyMorphS reproduces diverse axonal morphologies across WM regions

To assess whether interaction-driven mesh deformation produces realistic morphology at the population level, we evaluate PhyMorphS statistically against real 3D XNH segmentations containing thousands of axons from four WM regions (Amirrashedi et al., 2025). Rather than attempting to reproduce individual segmented axons, we compare distributions of diameter, longitudinal diameter variation, cross-sectional shape, tortuosity, and packing density. This evaluates whether anatomically realistic distributions can emerge from regional statistical priors combined with local physical interactions.

The synthesized ADDs agree with the XNH measurements over the reliably segmented diameter range. Because small axons are underrepresented in the XNH data, the underlying ADD is estimated from the truncated measurements rather than sampled directly from the observed histogram. The resulting initialization therefore includes sub-micrometer axons consistent with previous histological observations (Tian et al., 2025; Sepehrband et al., 2016; Lee et al., 2020b; Caminiti et al., 2013). During synthesis, the ADD shifts toward smaller diameters because mechanical and packing constraints prevent some axons from reaching their prescribed target radii. This demonstrates that the target radii should be interpreted as growth targets rather than guaranteed final dimensions.

Convexity and longitudinal diameter variation show relatively close agreement with the XNH data, whereas eccentricity exhibits a larger difference. The controlled experiment in Fig. 4.B indicates that eccentricity depends strongly on the initialized fiber arrangement in addition to interaction-driven deformation. Tortuosity shows a similar dependence on initialization: XNH axons in the splenium and midbody are predominantly straight, whereas CF and U-fibers exhibit broader tortuosity distributions. The synthesized substrates preserve this regional ordering but show less separation between these two organizational regimes.

Packing density further influences interaction-driven morphology. With the same target FVF of 80%, the achieved FVF varies from approximately 66% to 79% across regions, showing that orientation and packing configuration constrain the attainable axonal size. The relatively high target FVF was chosen to probe dense packing and strong local interactions. Real WM FVF is generally lower, and the morphology generated at lower packing densities may consequently exhibit weaker interaction-driven deformation. Conversely, the XNH reference tissue may exhibit comparatively high packing because of sample preparation and dehydration. Comparison with three-dimensional references acquired under conditions closer to native tissue, such as cryo-EM, would therefore provide an important means of studying how packing density affects morphology (Skoven et al., 2025).

Importantly, the evaluated metrics characterize complementary but limited aspects of morphology. Convexity measures deviation from a convex boundary, eccentricity quantifies anisotropy of the cross-sectional spread, and CoV*_d_* characterizes longitudinal diameter variation. The agreement in these statistics therefore does not imply complete geometric equivalence. Direct mesh-based representation in PhyMorphS is useful because it retains morphological information beyond any individual summary statistic. Thus, PhyMorphS can be used not only to reproduce measured morphological distributions but also to systematically investigate how packing density and local interactions reshape those distributions.

### 4.2. Contributors to morphological variation beyond local interactions

Our results indicate that the final axonal morphology is shaped by two complementary factors: the initial organization of the tissue and the subsequent deformation produced by local interactions. In the present experiments, axonal positions are sampled randomly and orientations are drawn independently from the regional FODs. The synthesized substrates therefore reproduce regional orientation and diameter statistics but are not intended to replicate the exact spatial organization of the corresponding XNH tissue.

This distinction is particularly relevant for eccentricity and tortuosity. Axonal trajectories depend on how fibers are routed through the tissue, while cross-sectional eccentricity is influenced by anisotropic packing and local density. The controlled experiment in Fig. 4.B supports this separation: changing the initialized fiber arrangement alters the resulting eccentricity while the PhyMorphS deformation model remains unchanged. Thus, the final morphology reflects both the organization prescribed at initialization and the interaction-driven deformation that develops during synthesis.

White matter organization is known to extend beyond local orientation statistics. For example, laminar and bundle-level arrangements have been observed across anatomical scales (Kjer et al., 2025), and previous synthesizers such as WMG have demonstrated that structured fiber configurations can be explicitly introduced during initialization (Winther et al., 2024b). In PhyMorphS, initialization is intentionally decoupled from the deformation pipeline. More anatomically informed centerlines, including laminar, bundle-level, or tractography-derived configurations, can therefore be supplied through the same configuration interface while retaining the same physics-guided growth and interaction model.

The framework can also incorporate additional structures that contribute to the local microstructural environment. Blood vessels and larger arrangements of cell bodies can alter local packing, axonal trajectories, and cross-sectional morphology (Andersson et al., 2020). Because collision handling operates on triangle meshes, such structures can be introduced as additional interacting objects without changing the underlying axonal representation. The same mesh-based formulation also provides a basis for progressively introducing more detailed cellular or intracellular components when their geometric influence is of interest (Kevenaar and Hoogenraad, 2015).

This separation between anatomical initialization and physics-guided deformation allows PhyMorphS to combine increasingly realistic tissue organization with local interaction-driven morphology. Rather than prescribing every geometric detail directly, the framework can use anatomical priors to define the larger-scale environment and allow the final local morphology to emerge through mechanical interactions within that environment.

### 4.3. Flexible synthesis of morphological alterations

PhyMorphS also provides direct control over prescribed morphological alterations while preserving interactions with the surrounding tissue. Axonal beading is generated through repeated swellings whose amplitude and frequency are independently controlled. Variations in axonal diameter and beading have been shown to influence diffusion measurements (Budde and Frank, 2010; Lee et al., 2020b), making controlled manipulation of these features useful for investigating their effect on simulated dMRI signals.

Because beading is introduced through the mass-spring system rather than imposed as a fixed final geometry, neighboring structures can modify the prescribed swelling pattern during synthesis. The same principle is not restricted to sinusoidal beading: arbitrary longitudinal target-radius profiles can be supplied to represent localized swelling or more complex diameter variations.

The multilayer mesh representation provides similar flexibility for myelin morphology. Myelin layer decompaction is represented by increasing interlamellar spacing while retaining irregular cross-sectional geometry. The representation can therefore support controlled variation in layer number, spacing, sheath thickness, and local geometry. Such flexibility is relevant because myelin morphology is closely linked to axonal structure and function and is increasingly considered in quantitative imaging models (Canales-Rodríguez et al., 2024; Skoven et al., 2025).

These examples are not intended to assign the synthesized shapes to a specific pathological mechanism. Rather, they demonstrate that PhyMorphS provides a controllable geometric platform in which axonal and myelin alterations can be varied independently and subsequently tested in downstream simulations.

### 4.4. Applications and practical considerations

The present work deliberately focuses on generating and validating the microstructural substrate itself rather than on a specific downstream dMRI simulation. PhyMorphS is intended to provide known geometric ground truth that can subsequently be used for Monte Carlo diffusion simulations. The resulting synthetic signals can then be analyzed with biophysical models, allowing model-derived estimates to be compared directly with the known microstructural properties of the substrate.

Supporting such downstream applications requires balancing geometric fidelity with computational and storage costs. Compared with primitive-based synthesizers such as MEDUSA (Ginsburger et al., 2019), the explicit triangle-mesh representation requires more memory and more detailed collision handling, but it also provides the geometric freedom needed to represent nonconvex, asymmetric, and locally varying morphologies. A typical (50 µm)^3^ substrate requires approximately 400 MB without myelin and around 1 GB when five myelin layers are represented. When the full geometric resolution is unnecessary, adaptive mesh resolution or mesh simplification could reduce this cost while preserving the morphological features relevant to the intended simulation.

A related trade-off arises from the synthesis objective itself. Existing frameworks adopt different strategies for achieving dense axonal packing (Ginsburger et al., 2019; Villarreal-Haro et al., 2023; Winther et al., 2024b). PhyMorphS prioritizes the sampled ADD by terminating growth according to the prescribed growth iterations and target radii. Local packing constraints can therefore prevent some axons from reaching their targets, resulting in a final FVF below the initialization target. In the current implementation, the target FVF is primarily used to estimate the number of axons required during initialization. Alternatively, FVF could be incorporated directly into the termination criterion when packing density is the primary objective, at the cost of modifying the resulting ADD. The growth criterion can therefore be adapted to emphasize either diameter fidelity or packing density depending on the intended application.

Spatial scale introduces a further consideration. The current morphological validation uses (50 µm)^3^ substrates and therefore focuses on local microstructure, whereas larger-scale organizational features, including the laminar arrangements described by Kjer et al. (2025), require structured centerlines extending beyond this scale. Because initialization is decoupled from the deformation pipeline, such organizational priors can be incorporated without redesigning the synthesis framework. Moreover, the approximately linear runtime scaling observed with increasing axon count provides a practical basis for extending PhyMorphS to larger substrates, while GPU acceleration and more efficient collision handling could further increase the accessible scale.

These properties make PhyMorphS suitable not only for generating individual realistic substrates but also for systematic model-development workflows. Its explicit closed-mesh representation can be directly used in Monte Carlo diffusion simulations to investigate how cross-sectional shape, diameter variation, packing density, myelin morphology, and tissue organization affect simulated dMRI signals. Repeating this process across independently synthesized substrates can generate large paired datasets of known microstructure and simulated signals. Such datasets provide controlled ground truth for evaluating conventional biophysical models and can also support the training of data-driven or learning-based models that infer microstructural properties from dMRI. In this way, PhyMorphS provides a common framework for substrate generation, model validation, and model development.

### 4.5. Reproducibility and data availability

PhyMorphS is not a machine-learning method. For a fixed configuration file, the synthesis algorithm is deterministic apart from the stochastic generation of multiple myelin layers, which can be controlled using a user-specified random seed. Identical configurations, parameter settings, and seeds therefore reproduce the synthesis results, subject to minor numerical differences arising from multithreaded execution and floating-point rounding.

We use all available data from the real XNH segmentation datasets across the four WM regions. These data provide regional statistics for initialization and anatomical references for morphological evaluation. Portions of the source data are publicly available through Andersson et al. (2020) and Kjer et al. (2025). The remaining data will be made publicly available upon publication of the corresponding studies.

## 5. Acknowledgement

RC was supported by a DTU-EPFL alliance scholarship. MA, MCB, and TBD were supported by the European Research Council (ERC) under the European Union’s Horizon Europe research and innovation programme (grant agreement No. 101044180, “CoM-BrainN”, PI: Tim B. Dyrby).

## 6. Declaration of generative AI and AI-assisted technologies in the manuscript preparation process

During the preparation of this work the author(s) used ChatGPT to improve the manuscript’s structure, clarity, and language. After using this tool/service, the author(s) reviewed and edited the content as needed and take(s) full responsibility for the content of the published article.

## Appendix A. FOD fitting of four WM regions

**Figure A.7:**
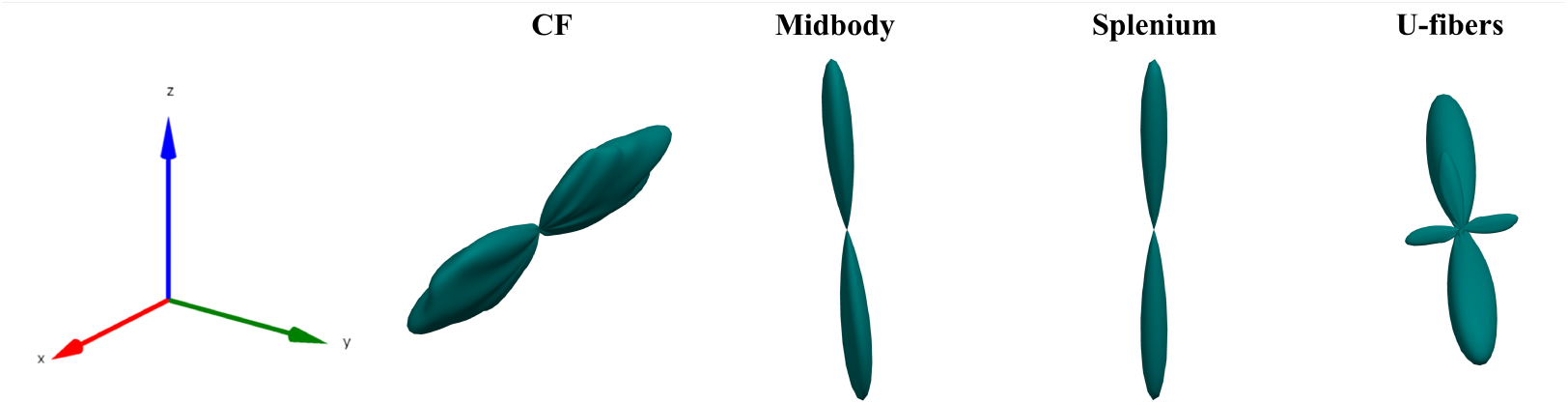
Fiber orientation distributions (FODs) estimated from the XNH data for the four WM regions: crossing fibers (CF), midbody, splenium, and U-fibers. The FODs are represented using spherical harmonics and visualized in a common coordinate system, with the reference axes shown on the left.

## Appendix B. Possible failure in dynamic simulation

Stable post-growth deformation requires the displacement per time step to remain sufficiently small relative to the dimensions of the interacting structures. Numerical instability can otherwise occur, particularly for thin axons subjected to large collision-induced displacements. Fig. B.8 presents one such example.

**Figure B.8:**
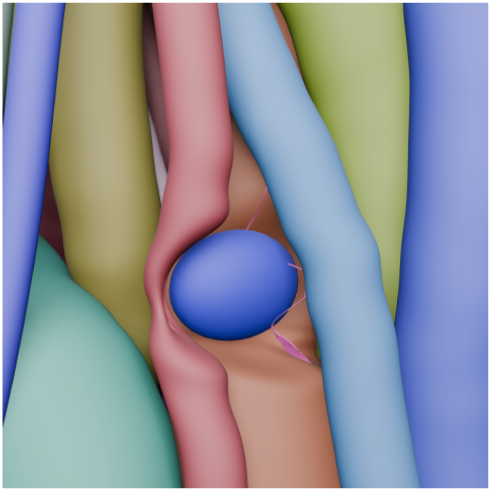
Failure case of the dynamic system simulation. The mesh of the thin axon in pink becomes unstable and eventually explodes after optimization from a collision-induced deformed state.

## Notes

### Competing Interest Statement

The authors have declared no competing interest.

